# Root Anatomy based on Root Cross-Section Image Analysis with Deep Learning

**DOI:** 10.1101/442244

**Authors:** Chaoxin Wang, Xukun Li, Doina Caragea, Raju Bheemanahallia, S.V. Krishna Jagadish

## Abstract

Aboveground plant efficiency has improved significantly in recent years, and the improvement has led to a steady increase in global food production. The improvement of belowground plant efficiency has potential to further increase food production. However, belowground plant roots are harder to study, due to inherent challenges presented by root phenotyping. Several tools for identifying root anatomical features in root cross-section images have been proposed. However, the existing tools are not fully automated and require significant human effort to produce accurate results. To address this limitation, we use a fully automated approach, specifically, the Faster Region-based Convolutional Neural Network (Faster R-CNN), to identify anatomical traits in root cross-section images. By training Faster R-CNN models on root cross-section images, we can detect objects such as root, stele and late metaxylem, and predict rectangular bounding boxes around such objects. Subsequently, the bounding boxes can be used to estimate the root diameter, stele diameter, late metaxylem number, and average diameter. Experimental evaluation using standard object detection metrics, such as intersection-over-union and mean average precision, has shown that the Faster R-CNN models trained on rice root cross-section images can accurately detect root, stele and late metaxylem objects. Furthermore, the results have shown that the measurements estimated based on predicted bounding boxes have small root mean square error when compared with the corresponding ground truth values, suggesting that Faster R-CNN can be used to accurately detect anatomical features. A webserver for performing root anatomy using the Faster R-CNN models trained on rice images is available at https://rootanatomy.org, together with a link to a GitHub repository that contains a copy of the Faster R-CNN code. The labeled images used for training and evaluating the Faster R-CNN models are also available from the GitHub repository.

## 1. Introduction

The crop scientific community has made significant strides in increasing global food production through advances in genetics and management, with majority of the progress achieved by improving aboveground plant efficiency [1, 2, 3]. The belowground plant roots, which provide water and nutrients for plant growth, are relatively less investigated. This is primarily because of the difficulty in accessing the roots, and the complexity of phenotyping root biology and function [4, 5]. Hence, root potential has largely been untapped in crop improvement programs [4, 5]. Over the past decade, different root phenotyping approaches have been developed for studying root architecture, including basket method for root angle [6], rhizotron method for tracking root branching, architecture and growth dynamics [7], shovelomics, a.k.a., root crown phenotyping [8], among others. Recent advances in magnetic resonance imaging and X-ray computed tomography detection systems have provided the opportunity to investigate root growth dynamics in intact plants at high temporal frequency [9, 10, 11, 12, 13]. However, each of these techniques comes with a range of inherent biases or limitations (such as artificial plant growth conditions), with none of the techniques currently available clearly standing out as a promising “blanket fit” approach [14, 15, 16]. Recent non-destructive technologies, such as X-ray computed tomography, are extremely expensive, and thus beyond the reach of common crop improvement programs, in addition to not having the bandwidth to capture large genetic diversity.

Machine learning is an area of artificial intelligence, focused on models that can automatically infer patterns from existing data, without human intervention [17]. In supervised machine learning, data in the form of training instances are provided as input, and the models learn patterns that can be used to make predictions on new unseen data. Machine learning approaches have been used successfully to address a wide variety of bioinformatics and computational biology problems relevant to crop sciences, including prediction of gene functions in plants [18], discovery of single nucleotide polymorphisms (SNP) in plyploid plants [19], subcellular localization [20], genomic selection for plant breeding [21], high-throughput plant phenotyping based on image analysis [22], prediction of biomass [23]. Furthermore, applications of advanced deep neural networks to challenging problems in crop analysis have led to state-of-the-art results that outperform the results of traditional machine learning and image analysis techniques approaches as reviewed in [24].

Most relevant to this work, machine learning, in general, and deep neural networks (a.k.a., deep learning), in particular, are expanding the ability to accurately predict a plant phenotype [25, 26, 27, 28, 29, 30, 31, 32, 33]. These technological advances have enabled researchers to capture a wide range of genetic diversity, a task which has been hardly possible in the past, given the amount of time and effort involved in manual analysis. Several recent studies have used deep learning approaches for identifying and quantifying aboveground plant traits, such as the number of leaves in rosette plants, based on high-resolution RBF images [29, 30, 31]. Other investigations have focused on identifying plant diseases [34] or on stress phenotyping [26].

Furthermore, several prior studies have focused on data-driven approaches and tools for belowground plant phenotyping, including identifying and quantifying root morphological parameters, such as changes in root architecture, or branching and growth [35, 36, 37, 38]. Such approaches rely on standard image analysis techniques as opposed to state-of-the-art deep learning.

Both root morphological and anatomical traits are important in relation to the efficiency of soil moisture absorption by the root system. Large genetic variation in root related traits has positioned rice to uptake water and increase yields under a range of ecological conditions, including flooded and dryland conditions [39]. Root anatomical traits such as nodal root diameter (RD) [40], late metaxylem diameter (LMXD) and number (LMXN) [41, 42, 43], and stele diameter (SD) and its proportion to root diameter (SD:RD) [44] have been proposed as key traits for optimized acquisition of water and productivity under water-limited conditions [40]. Thin SD:RD has been used as a surrogate measure of cortex tissue area/width, which helps in the improvement of water flow and retention in vascular tissue [45, 44]. Late metaxylem number and diameter along the root influence the hydraulic conductivity [41, 44]. These parameters mentioned above help to determine effective water use throughout the crop growth period [46, 43].

Innovations in image acquisition technologies have made it possible to gather relatively large sets of root cross-section images, enabling studies on root anatomy. Several approaches and tools for quantifying root anatomical variation based on cross-section images have been proposed in recent years [47, 48, 49]. However, the existing tools are only partially automated, as they require user input and fine-tuning of the parameters for each specific image or for a batch of images. Fully automated tools exist for the analysis of hypocotyl cross-sections (i.e., the region in between seed leaves and roots) in the context of secondary growth [50, 51], but they are not directly applicable to the analysis of root cross-section images. Thus, there is a pressing need for automated root cross-section image analysis tools that can be used to perform root anatomy at a low cost.

To address this limitation, we have taken advantage of recent advances in deep learning and image analysis, and used a state-of-the-art, fully-automated deep learning approach, the Faster R-CNN network [52], to identify and quantify root anatomical parameters indicative of physiological and genetic responses of root anatomical plasticity in field crops. Specifically, as a proof-of-concept, we have focused on the following parameters: root diameter (RD), stele diameter (SD), late metaxylem diameter (LMXD) and late metaxylem number (LMXN), which were found important in relation to water-deficit stress in our prior work [44, 53]. A graphical illustration of these parameters is shown in Figure 1.

**Figure 1:**
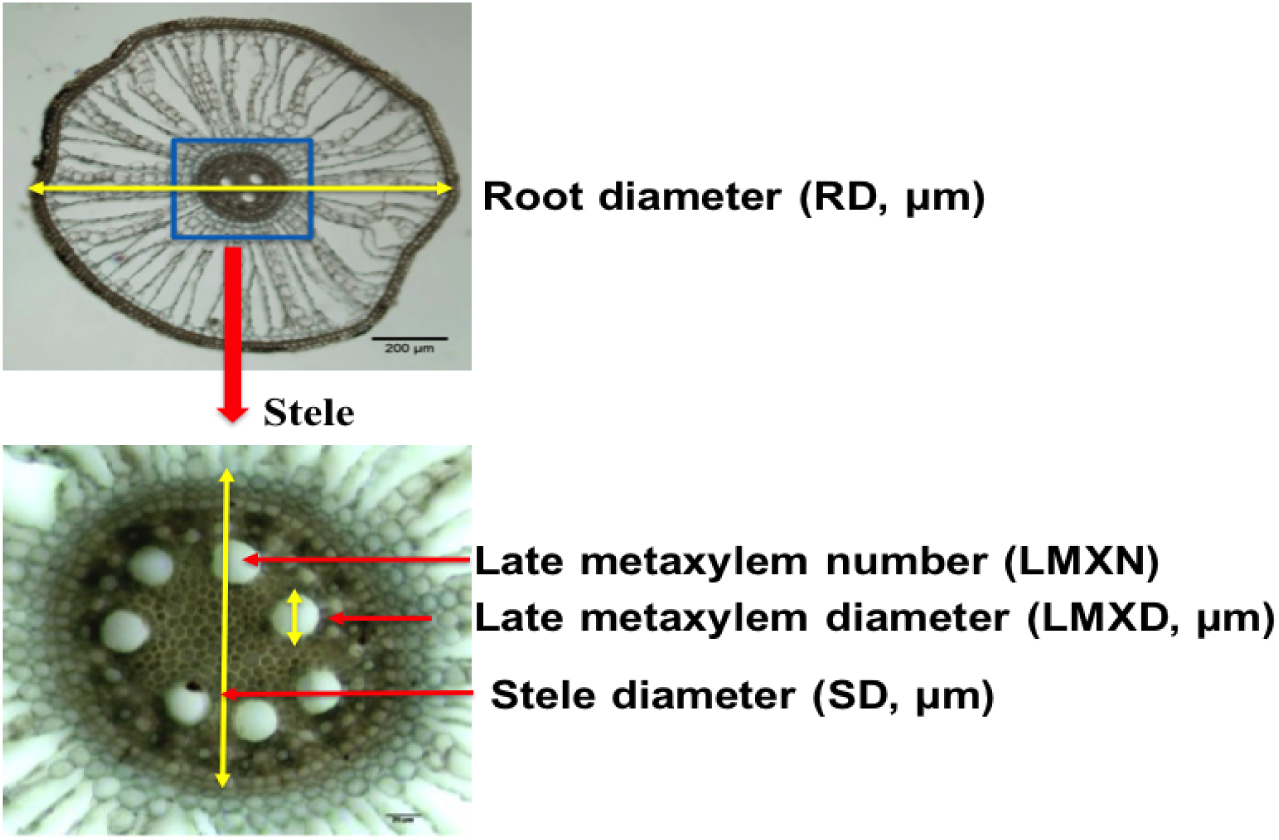
Root anatomical traits. (Top) Root cross-section with highlighted *root diameter* and *stele*. Image taken at 50x magnification. (Bottom) Enlarged stele with highlighted *stele diameter*, and *late metaxylem diameter*. The *late metaxylem number* is also a trait of interest. The image was taken at 100x magnification.

The existing Faster R-CNN model was trained on rice root cross-section images. The trained model was used to detect objects of interest in a root cross-section image (i.e., root, stele and late metaxylem), together with their corresponding bounding boxes. Subsequently, the bounding boxes were used to estimate anatomical parameters such as RD, SD, LMXD, LMXN. The Faster R-CNN model generalizes well to unseen images, thus eliminating the need for the end-user to hand-draw a stele border or manually choose or correct the metaxylem cells, tasks that are time-consuming, and also prone to noise and errors.

To summarize, our main contributions are as follows:

- We have used the Faster R-CNN network trained on root cross-section images to detect root, stele and late metaxylem objects, and their corresponding bounding boxes.
- We have investigated the Faster R-CNN model with respect to the number of instances needed to accurately detect objects of interest, and their corresponding bounding boxes.
- We have evaluated the ability of the predicted bounding boxes to produce accurate estimates for anatomical properties, and performed error analysis to identify sources of errors.
- We have identified advantages and disadvantages of Faster R-CNN approach for root anatomy by comparison with existing approaches for this task.

## 2. Materials and Methods

While there are many anatomical traits that can be identified, and measured or counted (e.g., RootScan outputs more than 20 anatomical parameters), as a proof-of-concept, we have focused on measuring the root diameter (RD), stele diameter (SD), and late metaxylem diameter (LMXD), and counting the number of late metaxylem inside the stele (LMXN). Our choice was motivated by studies by Kadam et al. [44, 53], who showed the importance of these traits in relation to water-deficit stress, and provided the ground truth dataset for our study. The tasks that we target can be achieved with modern object detection techniques, such as Faster R-CNN, as described below. In addition to the traits of interest (RD, SD, LMXD and LMXN), other traits can be estimated based on the objects detected with our trained Faster R-CNN models (e.g., stele area, average area of the late metaxylem). Furthermore, Faster R-CNN or Mask R-CNN models [54] can be trained to detect other objects, such as aerenchym and protoxylem objects, and their parameters, if data annotated with such objects becomes available.

### 2.1. Overview of the Approach

We have used Faster R-CNN [52], a state-of-the-art network for object detection, to detect objects of interest (i.e., root, stele, late metaxylem), and subsequently mark each object with a bounding box. More precisely, we have trained a Faster R-CNN model to identify the root and stele within a cross-section image, and another Faster R-CNN model to identify the late metaxylem within the stele region of a cross-section. Given the bounding box of an object, identified by the Faster R-CNN models trained on root cross-section images, we have calculated its diameter by averaging the width and height of the bounding box. The count of late metaxylem was obtained by counting the number of late metaxylem objects detected by the Faster R-CNN network.

The Faster R-CNN model architecture is shown in Figure 2. As can be seen, the Faster R-CNN has two main components. The first component consists of a Region Proposal Network (RPN), which identifies Regions of Interest (i.e., regions that may contain objects of interest), and also their location. The second component consists of a Fast R-CNN [56], which classifies the identified regions (i.e., objects) into different classes (e.g., root and stele), and also refines the location parameters to generate an accurate bounding box for each detected object. The two components share the convolutional layers of the VGG-16 network [55], which is used as the backbone of the Faster R-CNN model. More details on convolutional neural networks, VGG-16 and Faster R-CNN approach, which we used to detect objects and generate bounding boxes, are provided below.

**Figure 2:**
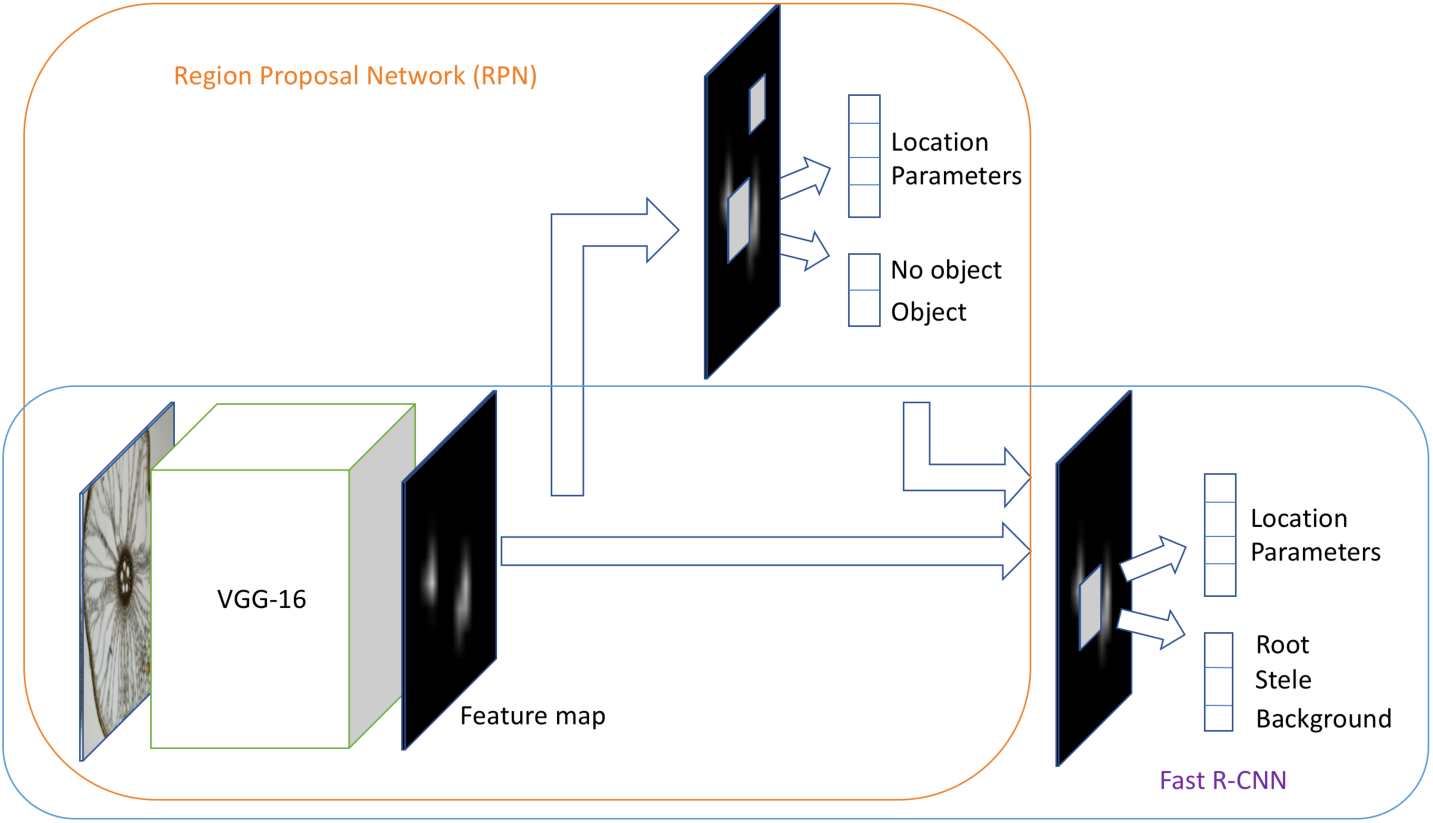
Faster R-CNN model architecture [52], which has two main components: 1) a region proposal network (RPN), which identifies regions that may contain objects of interest and their approximate location; and 2) a Fast R-CNN network, which classifies objects as root or stele, and refines their location, defined using bounding boxes. The two components share the convolutional layers of the pre-trained VGG-16 [55].

#### 2.1.1. Convolutional Neural Networks and VGG-16

Convolutional Neural Networks (CNNs) [57] are widely used in image analysis. While originally designed for image classification, the features extracted by CNNs are informative for other image analysis tasks, including object detection. A CNN consists of convolutional layers followed by non-linear activations, pooling layers and fully connected layers, as seen in Figure 3 (which shows a specific CNN architecture called VGG-16 [55]).

**Figure 3:**
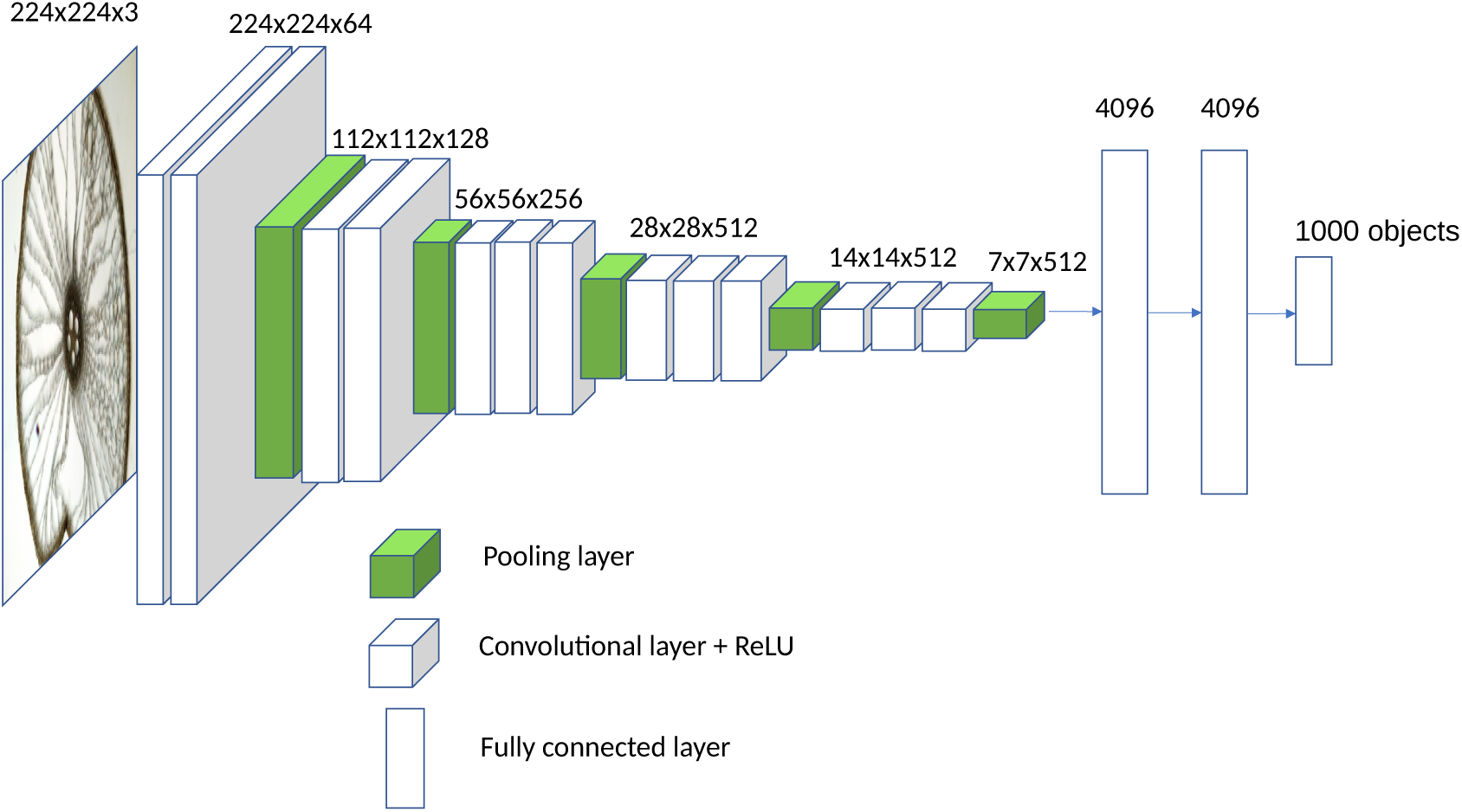
VGG-16. The original VGG-16 architecture consists of 13 convolution+ReLU layers, five pooling layers, and three fully connected layers. A convolution+ReLU layer produces a feature map, while a pooling layer reduces the dimensionality of the feature map. The last fully connected layer uses a softmax activation function to predict one of the 1000 categories. The dimensions corresponding to each layer are also shown.

A convolutional layer employs a sliding window approach to apply a set of filters (low-dimensional tensors) to the input image. The convolution operation captures local dependencies in the original image, and it produces a feature map. Different filters produce different feature maps, consisting of different features of the original image (e.g., edges, corners, etc.). A convolution layer is generally followed by a non-linear activation function, such as the Rectified Linear Unit (i.e., ReLU), applied element-wise to generate a rectified feature map. The ReLU activation replaces all the negative pixels in a feature map with zero values. A pooling layer is used to reduce the dimensionality of the rectified feature map. Intuitively, the pooling operation retains the most important information in a feature map by taking the maximum or the average pixel in each local neighborhood of the feature map. As a consequence, the feature map becomes equivariant to scale and translation [58].

After a sequence of convolutional layers (together with non-linear activations) and pooling layers, a CNN has one or more fully connected layers. In a fully connected layer, all neurons are connected to all neurons in the subsequent layer. The first fully connected layer is connected to the last downsized feature map. The fully connected layers are used to further reduce the dimensionality and to capture non-linear dependencies between features [58]. The last fully connected layer uses a softmax activation function, and has as many output neurons as the number of targeted classes.

There are several pre-trained CNN architectures available, including VGG-16 [55], shown in Figure 3. The VGG-16 network has been shown to give very good performance in the ImageNet competition, where the network was trained on millions on images with 1000 categories [55]. Furthermore, VGG-16 was used with good results in the original Faster R-CNN study [52], which motivated us to use it also in our study. As can be seen in Figure 3, VGG-16 has 13 convolutional+ReLU layers, 5 pooling layers, and 3 fully connected layers. The dimensions corresponding to each layer are also shown in Figure 3.

#### 2.1.2. Region Proposal Network (RPN)

As mentioned above, the region proposal network identifies regions that could potentially contain objects of interest, based on the last feature map of the pre-trained convolutional neural network that is part of the model, in our case VGG-16 [55]. More specifically, using a sliding window approach, *k* regions are generated for each location in the feature map. These regions, are represented as boxes called *anchors*. The anchors are all centered in the middle of their corresponding sliding window, and differ in terms of scale and aspect ratio [52], to cover a wide variety of objects. The region proposal network is trained to classify an anchor (represented as a lower-dimensional vector) as containing an object of interest or not (i.e., it outputs an “objectness” score), and also to approximate the four coordinates of the object (a.k.a., location parameters). The ground truth used to train the model consists of bounding boxes provided by human annotators. If an anchor has high overlap with a ground truth bounding box, then it is likely that the anchor box includes an object of interest, and it is labeled as positive with respect to the *object* versus *no object* classification task. Similarly, if an anchor has small overlap with a ground truth bounding box, it is labeled as negative. Anchors that don’t have high or small overlap with a ground truth bounding box are not used to train the model. During training, the positive and negative anchors are passed as input to two fully connected layers corresponding to the classification of anchors as containing *object* or *no object*, and to the regression of location parameters (i.e., four bounding box coordinates), respectively. Corresponding to the *k* anchors from a location, the RPN network outputs 2*k* scores and 4*k* coordinates.

#### 2.1.3. Fast R-CNN

Anchors for which the RPN network predicts high “objectness” scores are passed to the last two layers (corresponding to object classification and location parameter refinement, respectively) of a network that resembles the original Fast R-CNN network [56], except for how the proposed regions are generated. Specifically, in the original Fast R-CNN, the regions were generated from the original image using an external region proposal method (e.g., selective search).

As opposed to the original Fast R-CNN [56], in the Fast R-CNN component of the Faster R-CNN model, the external region proposal method is replaced by an internal RPN trained to identify regions of interest [52]. Highly overlapping regions, potentially corresponding to the same object, can be filtered using a non-maximum suppression (NMS) threshold. A pooling layer is used to extract feature vectors of fixed length for the regions of the interest proposed by RPN. Subsequently, the feature vectors are provided as input to two fully connected layers, corresponding to the classification of the object detected and the regression of its location, respectively.

The object classification layer in Fast R-CNN uses the softmax activation, while the location regression layer uses linear regression over the coordinates defining the location as a bounding box. All parameters of the network are trained together using a multi-task loss [56].

#### 2.1.4. Faster R-CNN Implementation and Training

The implementation of the original Faster R-CNN model [52], which is publicly available at https://github.com/ShaoqingRen/faster_rcnn, uses MAT-LAB as the programming language, and Caffe (http://caffe.berkeleyvision.org) as the backend deep learning framework. Chen and Gupta [59] provided an implementation of the Faster R-CNN model, which uses Python as the programming language and TensorFlow (https://www.tensorflow.org) as the backend deep learning framework. This implementation, publicly available at https://github.com/endernewton/tf-faster-rcnn, allows the user to train a model from scratch and also to reuse one of several pre-trained models as the backbone of the network. In particular, the user can select the VGG-16 network, pre-trained on the ImageNet dataset with 1000 categories.

We used the Python/TensorFlow implementation of the Faster R-CNN network, with the pre-trained VGG-16 model as its backbone, and trained the network to identify objects such as root, stele and late metaxylem. More precisely, the parameters of the VGG-16 convolutional layers, which are shared by the Fast R-CNN and RPN networks in Faster R-CNN, were initialized using the pre-trained VGG-16 network. As many image features are highly transferable between different datasets, this initialization based on VGG-16 allowed us to train accurate models from a relatively small number of root cross-section labeled images. In our preliminary experimentation, we found that it is difficult to accurately detect late metaxylem at the same time with root and stele. To address this issue, we trained a Faster R-CNN model to detect root and stele from background (i.e., everything else in the image), and another Faster R-CNN model to detect late metaxylem from background. To achieve this, we changed the output layer of the original Faster R-CNN network to reflect our classes (corresponding to the objects detected).

Given that the RPN and Fast R-CNN networks share 13 convolutional layers (initialized based on VGG-16), they were co-trained using an iterative process that alternates between fine-tuning the RPN and fine-tuning the Fast R-CNN network (with fixed proposed regions produced by RPN) [52]. All the model parameters were updated using stochastic gradient descent (SGD).

### 2.2. Existing Approaches for Root Anatomy

There are several approaches and tools for quantifying root anatomical variation based on cross-section images [47, 48, 49]. Approaches in this category can be roughly categorized as manual, semi-automated, and automated approaches. Manual analysis of root images relies heavily on subjective assessments, and is suitable only for low throughput analysis. ImageJ [60] is an image analysis tool that has been extensively used to manually identify and quantify root anatomical traits [44, 61, 53], given that it enables researchers to mark objects of interest and obtain their measurements. In particular, the ImageJ software was used to acquire the ground truth (in terms of quantitative annotations) for the images used in this study, specifically, RD, SD, LMXD and LMXN measurements.

Semi-automated tools require user feedback to tune parameters for individual images in order to get accurate results. *RootScan* [47] and *PHIV-RootCell* [49] are semi-automated tools that identify and quantify anatomical root traits. RootScan was originally designed for analyzing maize root cross-section images. The analysis of each image involves several steps. RootScan starts by isolating the cross-section from the background using a global thresholding technique [62]. Subsequently, the stele is segmented based on the contrast between pixel intensities within and outside the stele. Different cells within the stele (e.g., late metaxylem) are classified based on their area according to background knowledge on root anatomy for a particular species. RootScan can detect several types of objects (including lucunae, metaxylem and protoxylem), and also a broad range of parameters for each detected object. After each step, the user has to “approve” the automated detection or alternatively correct it, before moving to the next step. The tool can be run on a set of images in batch mode, but the user still needs to provide input for each step of the analysis for each image, as explained above.

The *PHIV-RootCell* tool for root anatomy is built using the ImageJ software [60], and provides options for selecting regions of interest (ROI) such as root, stele, xylem, and for measuring properties of these regions. It was designed for analyzing rice root cross-section images. Similar to RootScan, domain knowledge is used to identify ROIs. The PHIV-RootCell tool uploads and analyzes one image at a time, and does not have an option for batch uploading or processing. Furthermore, it requires user’s supervision at each segmentation and classification step [49]. For example, it requires the user to validate the root selection, stele selection, central metaxylem selection, among others.

As opposed to semi-automated tools that require user feedback, a fully automated approach should involve “a single click” and should produce accurate results without any human intervention during the testing and evaluation phases. However, human input and supervision in the form of background knowledge or labeled training examples may be provided during the training phase. In this sense, *RootAnalyzer* [48] is an automated tool, which incorporates background knowledge about root anatomy. The first step in RootAnalyzer is aimed at performing image segmentation to distinguish between root pixels (corresponding to boundaries of individual root cells) and background pixels. To achieve this, RootAnalyzer utilizes a local thresholding technique to analyze each pixel’s intensity by comparing it with the mean pixel intensity in a small square neighborhood around that pixel (defined by a width parameter, *W*). Subsequently, RootAnalyzer constructs a difference image, and classifies pixels as root or background pixels based on a threshold, *T*, used on the difference image. The next step is focused on detecting root cells and closing small leaks in cell boundaries, using an interpolation approach. Finally, cells are classified in different categories, such as stele cells, cortex cells, epidermal cells, etc. based on size, shape, and position. Two thresholds are used to classify cells as small or large: a threshold, *A_s_*, for small cells, and a threshold, *A_l_*, for large cells. Furthermore, stele cells are classified based on an additional threshold, *N*, on the maximum distance from a cell to any of its nearest neighbor cells. The RootAnalyzer tool can be used for both single image processing and batch processing. Single image processing allows the user to adjust and tune parameters, and also to interact with the tool at each stage of the segmentation and classification. Batch processing requires the user to provide the parameters to be used with a specific batch of plant images. Similar to RootScan, RootAnalyzer outputs a table of area measurements and counts for regions of interest. This tool was designed for wheat and was shown to work also for maize [48].

### 2.3. Dataset

Twenty-five accessions of Oryza species were grown in plastic pots (25 cm in height; 26 and 20 cm diameter at the top and bottom, respectively), filled with 6 kg of clay loam soil. Three replications per each accession were maintained under well-watered conditions and roots were sampled 60 days after sowing, to ensure fully mature roots. The roots were harvested and washed thoroughly. To obtain the cross-section images used in this study, root samples stored in 40% alcohol were hand sectioned with a razor blade using a dissection microscope. For each of the 25 rice accessions, and for each of the three biological replicates, root samples from root-shoot junction and 6 cm from the root tip were obtained. Images of root sections were acquired with the Axioplan 2 compound microscope (Zeiss, Germany) at 50x and 100x magnification. Specifically, for each accession and each replicate, 2-3 images were taken at root-shoot junction, and 2-3 images at 6 cm from the tip of the root, at 50x and 100x magnification. Thus, an image may have two versions: a 50*×* magnification version, which captures the whole root diameter (top image in Figure 1), and a 100*×* magnification version, which captures only the stele diameter (bottom image in Figure 1). However, not all 50*×* images have a 100*×* correspondent. Precisely, there are 388 images at 50*×* magnification, and 339 images at 100*×* magnification.

For each root image, we manually measured root anatomical parameters, such as root cross-section diameter, stele diameter, late metaxylem average diameter and late metaxylem number, using the ImageJ software [60]. Specifically, root diameters were estimated using the 50*×* magnification images. The stele diameter, and late metaxylem average diameter and count were estimated using the 100*×* magnification images, if available (otherwise, the 50*×* magnification images were used). The manual measurements and counts constitute our ground truth to which we compared the measurements produced based on the bounding boxes detected by our trained Faster R-CNN models. Statistics about the dataset, including the minimum, maximum, average and standard deviation for RD, SD, LMXD and LMXN, are presented in Table 1.

**Table 1:**
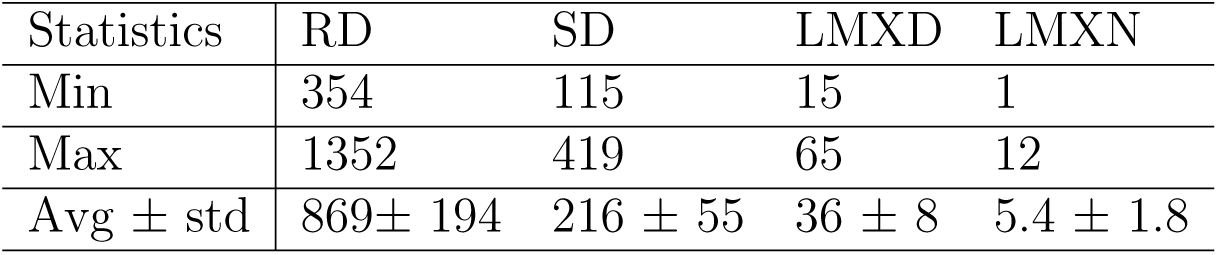
Ground Truth Statistics: minimum (Min), maximum (Max), and average together with standard deviation (Avg ± std) are shown for the ground truth measurements of RD, SD, LMXD (expressed in micrometers, *µm*) and LMXN (which is the count of late metaxylem objects).

| Statistics | RD | SD | LMXD | LMXN |
| --- | --- | --- | --- | --- |
| Min | 354 | 115 | 15 | 1 |
| Max | 1352 | 419 | 65 | 12 |
| Avg ± std | 869± 194 | 216 ± 55 | 36 ± 8 | 5.4 ± 1.8 |

In addition to measuring root anatomical parameters, each 50*×* magnification image was also manually labeled by independent annotators with bounding boxes that represent root, stele, and late metaxylem, respectively, and each 100*×* magnification image was labeled with boxes that represent late metaxylem.

We used the LabelImg tool available at ttps://github.com/tzutalin/labelImg to perform the bounding box labeling. This tool produces annotations in the Pascal Visual Object Classes (VOC) XML format [63], a standard format used for annotating images with rectangular bounding boxes corresponding to objects. An example of a root cross-section image annotated using the LabelImg tool is shown in Figure 4 (a), where each target object is marked using four coordinates, shown as green dots, which determine a bounding box. The bounding boxes annotated with the LabelImg tool in the 50*×* and 100*×* magnification images constitute the ground truth to which we compared the bounding boxes of the objects detected by our models. Corresponding to the ground truth image in Figure 4 (a) annotated with LabelImg, Figure 4 (b) shows the bounding box annotations produced by our models, as red boxes.

**Figure 4:**
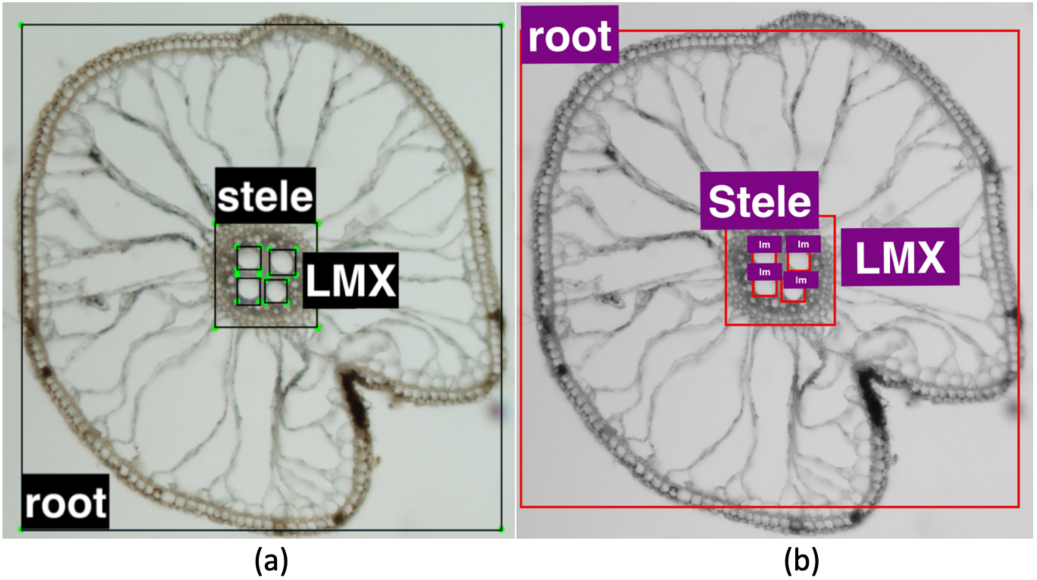
Objects of interests as bounding boxes: (a) Ground truth image annotated using LabelImg, where each object is marked using four coordinates, shown as green dots, which determine a bounding box. (b) The annotation of the same image by the root/stele and late metaxilem models, where the detected objects are shown using red bounding boxes.

We would like to emphasize that the 50*×* magnification images contain all the anatomical features that we target in this study, and are sufficient for training the proposed deep learning models. However, we also trained models on the 100*×* magnification images, independently, to understand how much the identification of the LMX objects and their measurements may be improved by using images with a higher resolution. In general, any resolution can be used for training, as long as all the features that need to be identified are contained in the image.

### 2.4. Experimental Setup

#### 2.4.1. Training, Development and Test Datasets

We performed a set of experiments using 5-fold cross-validation. Specifically, we split the set of 50*×* magnification images into five folds, based on accessions, such that each fold contained 5 accessions out of the 25 accessions available. The exact number of 50*×* magnification images (instances) in each fold is shown in Table 2. For each fold, Table 2 also shows the number of corresponding 100*×* magnification images (instances) available (as mentioned before, not every 50*×* magnification image has a corresponding 100*×* magnification image). In each 5-fold cross-validation experiment, four folds were used for training, and the fifth fold was used for test. To tune hyper-parameters, we used one of the training folds as the development dataset. The results reported represent averages over the 5 folds. The reason for splitting the set of images based on accessions was to avoid using images from the same plant or the same replicate both in the training and test datasets.

**Table 2:** Number of instances in each of the 5 folds used to perform cross-validation for the 50 and 100 magnification images, respectively. The total number of instances in the dataset is also shown.

| Fold | Fold 1 | Fold 2 | Fold 3 | Fold 4 | Fold 5 | Total |
| --- | --- | --- | --- | --- | --- | --- |
| Instances (50×) | 71 | 79 | 86 | 77 | 75 | 388 |
| Instances (100×) | 62 | 60 | 80 | 69 | 68 | 339 |

#### 2.4.2. Evaluation Metrics

We used three standard metrics in our evaluation, driven by preliminary observations. First, given that there exist exactly one root and one stele in an image, we observed that these objects are always detected in the 50*×* magnification images. We used the Intersection-over-Union (IoU) metric to measure how well the predicted bounding boxes overlap with the ground truth bounding boxes. Second, given that the number of LMX objects varies between 1 and 12, and these objects are relatively small, the corresponding object detection models are prone to both false positive and false negative mistakes. Thus, we used mean average precision (mAP), a standard metric in object detection, to evaluate the ability of our models to accurately identify the LMX objects. Both IoU and mAP metrics range between 0 and 1, and higher values are better. Finally, we used the root mean square error (RMSE) and relative root mean square error (rRMSE) (i.e., percentage error) metrics to measure the ability of the Faster R-CNN approach to detect objects and corresponding bounding boxes that lead to root/stele/LMX diameter measurements and LMX counts close to those available as ground truth. For RMSE and rRMSE, smaller values are better.

#### 2.4.3. Hyper-parameter Tuning

Deep learning models, in general, and the Faster R-CNN models, in particular, have many tunable hyper-parameters. We tuned several hyper-parameters shown to affect the performance of the Faster R-CNN models [64], and used the values suggested by Ren et al. [52] for the other hyper-parameters. More specifically, we tuned the IoU threshold used in the RPN network to identify anchors that could potentially include an object of interest (i.e., positive instances/anchors). Furthermore, we tuned the non-maximum suppression (NMS) threshold which is used to filter region proposals produced by the trained RPN network (specifically, if two proposals have IoU larger than the NMS threshold, the two proposals will be considered to represent the same object). At last, we tuned the fraction of positive instances in a mini-batch.

The specific values that we used to tune the IoU threshold were 0.4, 0.5 and 0.6; the values used to tune the NMS threshold were 0.6, 0.7 and 0.8; and the values used to tune the fraction of positive instances in a mini-batch were 1:5 and 1:4. To observe the variation of performance with the tuned parameters, and select the values that gave the best performance, we trained a model corresponding to a particular combination of parameters on three training folds, and evaluated the performance of the model on the development fold. The performance of the models for root and stele detection was measured using the IoU metric (by comparing the predicted bounding boxes with the ground truth bounding boxes), while the performance of the models for LMX detection was measured using the mAP metric (by comparing the detected LMX objects with the ground truth LMX objects) to ensure that the Faster R-CNN models can accurately detect all the LMX objects.

Our tuning process revealed that the performance did not vary significantly with the parameters for our object detection tasks. However, the best combination of parameters for the root/stele models consisted of the following values: 0.4 for the IoU threshold, 0.8 for the NMS threshold and 1:4 for the fraction of positive anchors in a mini-batch. The best combination of parameters for the LMX models was: 0.5 for the IoU threshold, 0.8 for the NMS threshold, and 1:4 for the fraction of positive anchors in a mini-batch. We used these combinations of values for the root/stele and LMX models, respectively, in our experiments described in the next section.

## 3. Results and Discussion

In this section, we present and discuss the results of our experiments using the Faster R-CNN models trained on rice root cross-section images. Furthermore, we outline time requirements for Faster R-CNN and discuss the availability of the Faster R-CNN model for root anatomy as a tool.

### 3.1. Variation of Performance with the Number of Training Instances

As opposed to the existing tools for identifying anatomical parameters in root cross-section images, which incorporate background knowledge about the root anatomy of a particular species and the types of images used, the automated Faster R-CNN approach is easily generalizable to various species and types of images, given that a representative set of annotated images is provided as training data. Under the assumption that data annotation is expensive and laborious, we aim to understand how many images are necessary for good performance on roots from a particular species. Intuitively, the number of required images should be relatively small, given that our model relies on a VGG-16 network pre-trained to detect a large number of objects, generally more complex than root, stele and late metaxylem objects.

To validate our intuition, we have performed an experiment where we varied the number of images used for training, while keeping the number of test images fixed. Specifically, we used 5, 10, 25, 50, 75, 100, 150, 200, 250, and all available training images in a split, respectively, to train models for detecting the root, stele and LMX in an image. The 50*×* magnification images were used to train the models for root/stele/LMX. The 100*×* magnification images were also used to train models for LMX, with the goal of understanding the benefits provided by higher resolution images. The trained models were subsequently used to detect root, stele, and LMX objects in test images.

The performance of the models was measured by comparing the predicted objects with the ground truth objects. We used the IoU metric to evaluate the predicted bounding boxes for root/stele by comparison with the corresponding ground truth bounding boxes. We used the mAP metric to measure the ability of the models to accurately detect LMX objects. The variation of performance with the number of training images is shown in Figure 5 for root/stele (Left plot) and LMX (Right plot). As can be seen, in the case of the models trained on the 50*×* magnification images, the performance increases with the number of training images, but tends to stabilize generally around 250 images. This confirms our intuition that only a small number of labeled images is needed to learn accurate models for the problem at hand. Furthermore, the left plot in the figure shows that the IoU values for both root and stele objects are around 0.95, when all the training images are used, and that the root bounding boxes are slightly better than the stele bounding boxes. Similarly, the LMX objects are detected with high accuracy, as shown in the right plot of Figure 5, where the mAP values are close to 0.9 consistently for models trained with smaller or larger number of 100*×* magnification images. Similar performance is obtained with the models trained from all 50*×* magnification images. The plots for both root/stele and LMX also show that generally the variance decreases with the size of the data. The slow decrease in performance that is observed sometimes between two training set sizes can be explained by the addition of some inconsistently labeled images present in the original dataset. Examples of inconsistently labeled images as shown in Figure 6.

**Figure 5:**
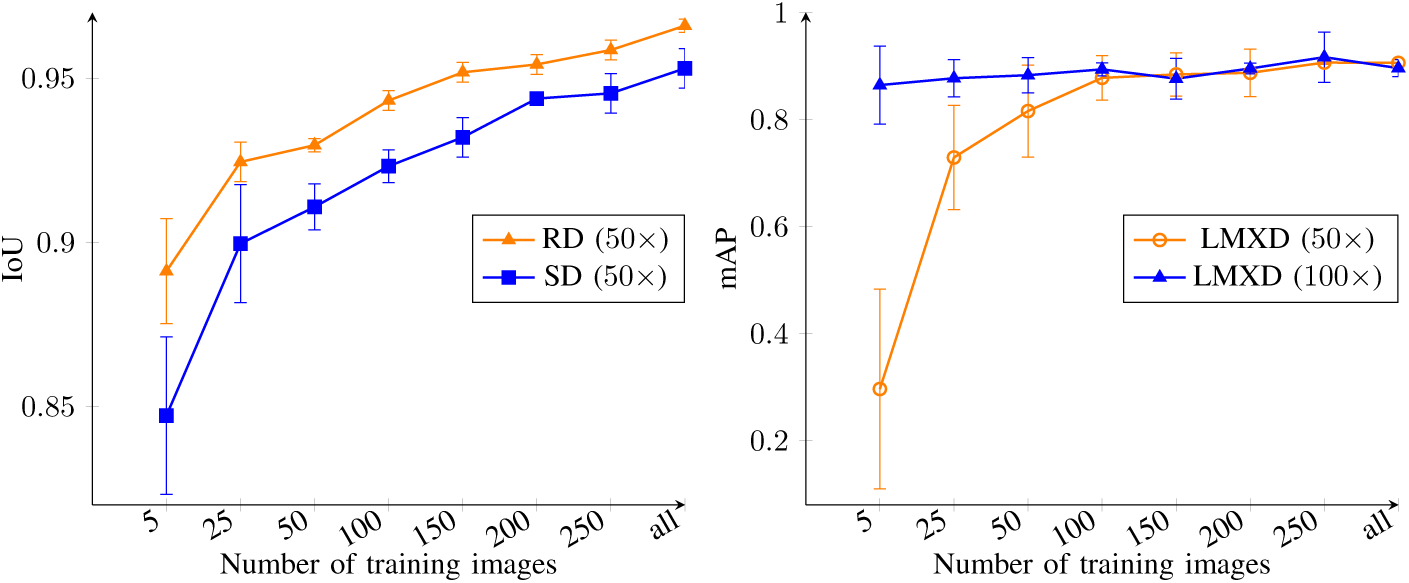
Variation of performance with the number of training images for root/stele detection model (Left plot), and for the LMX detection model (Right plot), respectively. We used 50 magnification images to detect root and stele objects, and both 50 and 100 magnification images to detect LMX. The performance of the root/stele detection model was measured using the IoU metric (which shows how accurately the predicted bounding boxes match the ground truth), while the performance of the LMX detection model was measured using the mAP metric (which shows how accurately LMX objects were detected). The plots show average values over 5 splits together with standard deviation.

**Figure 6:**
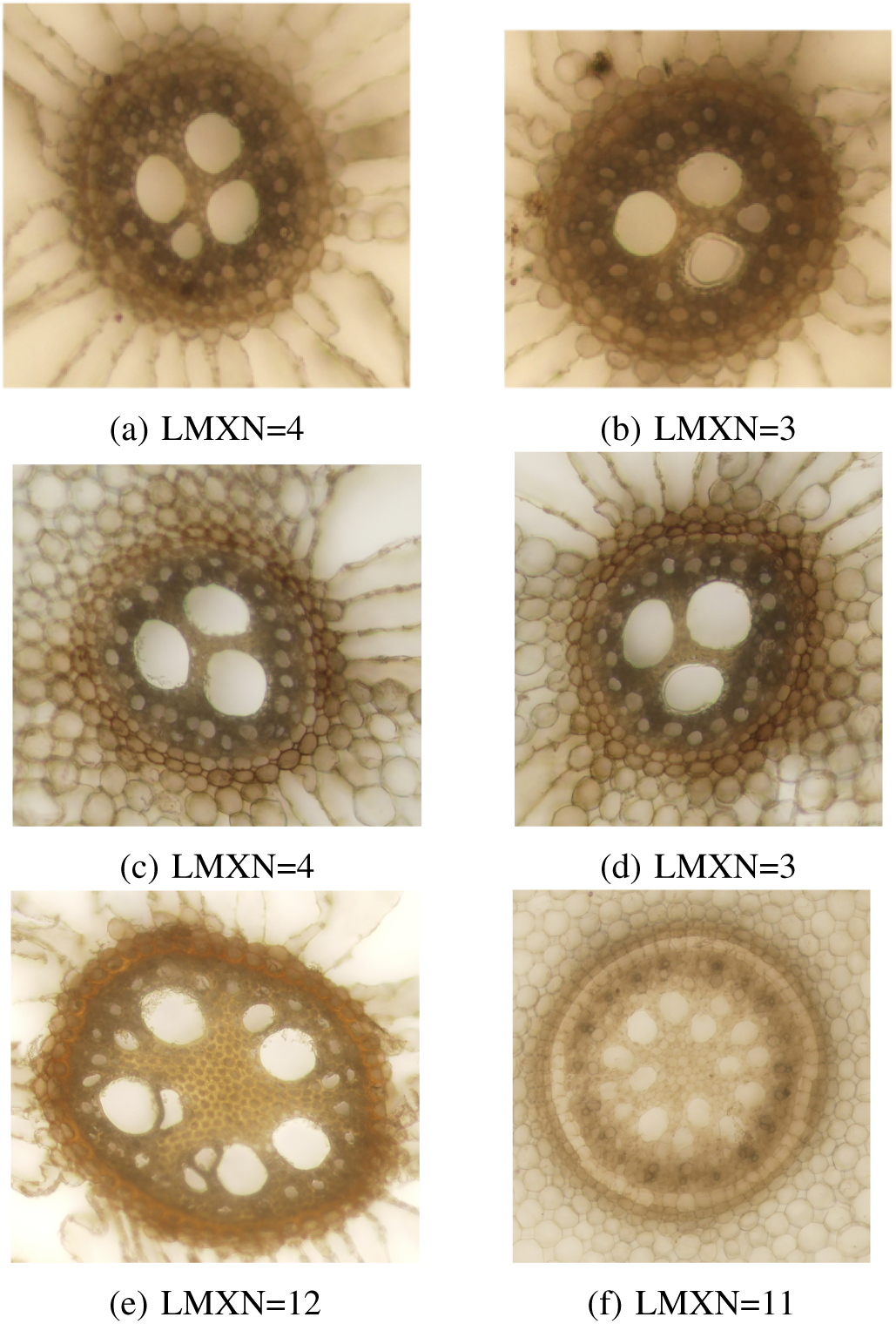
Examples of inconsistent human annotations that are included in our ground truth dataset. Specifically, image (a) was manually labeled as having LMXN=4 (the smaller LMX was included in the count), while image (b) was labeled as having LMXN=3 (the smaller LMX was not included in the count although it has size comparable with the smaller LMX counted in (a)). Our models consistently identified 4 LMX objects in both (a) and (b) images. Similarly, image (c) was incorrectly labeled manually as having LMXN=4, while the similar image in (d) was properly labeled as having LMXN=3. Our models correctly identified 3 LMX objects in both (c) and (d) images. Finally, images (e) and (f) show a larger number of LMX which have variable size, but it is not very clear which LMX were counted by the human annotator and which were not counted to get the 12 and 11 counts, respectively. Our models identified 7 LMX objects in image (e) and 10 LMX objects in image (f).

### 3.2. Performance Evaluation Using RMSE/rRMSE

The Faster R-CNN models trained on root images were used to detect root/stele/LMX objects in the test data. Subsequently, the detected objects were further used to calculate RD, SD, LMXD and LMXN. To evaluate the models in terms of their ability to produce accurate root/stele/LMX diameter and LMX number, we have used the RMSE error computed by comparing the measurement/count estimates obtained from the predicted bounding boxes with the ground truth measurements/counts. The RD and SD measurements were evaluated based on models trained/tested with the 50*×* magnification images, while LMXD and LMXN were evaluated based on models trained/tested with 50*×* and 100*×* magnification images, respectively. Intuitively, the LMXD/LMXN results obtained with the models trained on the 100*×* magnification images should be more accurate, as those images have higher resolution. The RMSE/rRMSE results of the experiments corresponding to the five splits, together with the average over the five splits, are shown in Table 3. In addition, Table 3 shows the expected human error estimated by performing an additional manual annotation using ImageJ (similar to how the original ground truth annotation was done), and comparing the second manual annotation against the first manual annotation.

**Table 3:** RMSE (*µm*) and rRMSE (i.e., percentage error) results for root diameter (RD), stele diameter (SD), late metaxylem diameter (LMXD) and late metaxylem number (LMXN) for 5 splits, together with the average over the 5 splits, and also the estimates for the human error. The number of 50x magnification images used in these experiments is 388, while the number of 100x magnification images is 339. For each measurement, the magnification of the images used to train the model that produced that measurement (i.e., 50*×* or 100*×*) is also shown.

| Split | RD($50\times$) | | SD($50\times$) | | LMXD($50\times$) | | LMXD($100\times$) | | LMXN($50\times$) | | LMXN ($100\times$) | |
| --- | --- | --- | --- | --- | --- | --- | --- | --- | --- | --- | --- | --- |
|  | RMSE | rRMSE | RMSE | rRMSE | RMSE | rRMSE | RMSE | rRMSE | RMSE | rRMSE | RMSE | rRMSE |
| Split 1 | 62.77 | 6.78 | 21.93 | 9.16 | 3.67 | 9.50 | 2.45 | 6.54 | 0.81 | 22.34 | 1.37 | 24.55 |
| Split 2 | 32.18 | 3.94 | 17.54 | 8.32 | 3.77 | 10.53 | 3.13 | 8.18 | 0.71 | 16.55 | 0.45 | 9.17 |
| Split 3 | 61.19 | 6.90 | 21.96 | 9.16 | 3.53 | 9.07 | 3.22 | 7.87 | 0.91 | 17.35 | 0.83 | 15.53 |
| Split 4 | 33.12 | 3.74 | 20.01 | 9.18 | 3.58 | 11.70 | 3.56 | 10.34 | 1.90 | 30.98 | 0.63 | 11.33 |
| Split 5 | 43.67 | 3.26 | 20.94 | 10.26 | 2.43 | 7.51 | 1.61 | 4.61 | 0.74 | 16.39 | 0.25 | 5.02 |
| Average | 46.59 | 4.92 | 20.39 | 9.21 | 3.40 | 9.66 | 2.79 | 7.51 | 1.02 | 20.72 | 0.71 | 13.12 |
| Human error | 48.14 | 5.46 | 25.17 | 11.29 | 3.39 | 9.13 | 3.39 | 9.13 | 0.21 | 3.89 | 0.21 | 3.89 |

As can be seen from Table 3, the average RMSE error for RD over the 5 splits is 46.59*µm*, while the average rRMSE is 4.92%. Given that root diameter for the images in our dataset varies between 354*µm* and 1352*µm* (see Table 1), and that the RMSE estimate for human error for RD is 48.14*µm* (with the corresponding rRMSE being 5.46%), these results suggest that the Faster R-CNN models trained on rice images can accurately learn to predict RD. Similarly, the average RMSE error for SD over the five splits is 20.39*µm* and the corresponding rRMSE is 9.21%, while the stele diameter varies between 115*µm* and 419*µm*. As for RD, the RMSE/rRMSE errors for the SD predictions are smaller than the estimates for human error, which are 25.17*µm* and 11.29%, respectively. As opposed to root and stele, the LMXD is significantly smaller, varying between 15*µm* and 65*µm*. In this case, the average RMSE error is 3.40*µm* and 2.79*µm* for models trained using 50*×* and 100*×* magnification images, respectively. The rRMSE for the model trained on the 50*×* magnification images is 9.66%, and decreases to 7.51% for the model trained on the 100*×* magnification images. Compared with the SD estimates for human error (which are based on the 100*×* magnification images, when available, or the 50*×* magnification images, otherwise), the results of the models trained on the 50*×* magnification images are slightly worse (rRMSE is 9.66% versus 9.13%), while the results of the models trained on the 100*×* magnification images are slightly better (7.51% versus 9.13%).

In terms of LMXN, the ground truth numbers vary between 1 and 12, with an average of 5 LMX objects per image. The average RMSE error for LMXN is 1.02 for models trained on 50*×* magnification images and 0.71 for models trained on 100*×* magnification images. Correspondingly, the rRMSE is 20.70% for models trained on 50*×* magnification images, and down to 13.12% for models trained on 100*×* magnification images. While the Faster R-CNN models trained with the 100*×* magnification images reduce the rRMSE error by approximately 7.5%, their average error is still higher than the estimate for human error by approximately 10%, showing that these models could be further improved with more training data.

We performed error analysis to gain insights into the usefulness of these results in practice. Specifically, we analyzed images where our models made mistakes in terms of LMXN, and observed that some of those images were annotated in an inconsistent way by the human annotators, as can be seen in Figure 6, where some smaller LMX objects are sometimes counted and other times not counted. This observation is not surprising, as human annotators are prone to mistakes and inconsistencies. As opposed to that, the automated Faster R-CNN models produce more consistent results (i.e., consistently count or not count a smaller LMX). More training images are necessary to learn well in the presence of noise/inconsistencies. Nevertheless, our results suggest that the Faster R-CNN approach to root anatomy has the potential to replace the labor-intensive manual annotations of root cross-section images.

### 3.3. Faster R-CNN Robustness to Image Variations

We further studied the ability of the Faster R-CNN models to “adapt” to other types of root cross-section images. To do this we identified 14 images that have been used to demonstrate RootAnalyzer and 10 images that have been used to demonstrate PHIV-RootCell. In addition, we also searched the Web for root cross-section images, and identified 15 more images from rice, 9 images from maize, and 9 images labeled as monocot root cross-section images. Together, our dataset of *external* images consists of 57 heterogeneous images, which came from different species, were taken with different imaging technologies under different conditions, had different sizes and resolutions, different backgrounds, different luminosity, etc. We randomly split each category of images into training/validation and test subsets. Specifically, 42 images were used for training/validation and 15 images were used for test. We initially used the Split 1 models (trained on 50*×* magnification images) to identify RD, SD, LMXD and LMXN traits for the external test images. Subsequently, we fine-tuned the Split 1 models with the external training images, and used the fine-tuned models to identify the RD, SD, LMXD and LMXN traits for the external test images. The results of these experiments are shown in Table 4.

**Table 4:** Faster R-CNN Model Robustness to Image Variations. The training and test internal images correspond to the training and test subsets of Split 1. The external images are collected from the Web. We used RMSE(*µm*)/rRMSE(%) to compare models trained on internal images with models trained on internal and external images in terms of their ability to detect RD/SD/LMX objects (and derived their diameter) in a variety of images.

| Experiment | RD (50%) |  | SD (50%) |  | LMXD (100%) |  | LMXN (100%) |  |
| --- | --- | --- | --- | --- | --- | --- | --- | --- |
|  | RMSE | rRMSE | RMSE | rRMSE | RMSE | rRMSE | RMSE | rRMSE |
| Train on internal images<br>Test on external images | 480.99 | 57.14 | 301.46 | 100.28 | 45.02 | 91.04 | 3.78 | 53.96 |
| Train on internal/external images<br>Test on external images | 24.85 | 2.95 | 13.67 | 4.55 | 3.85 | 7.79 | 0.58 | 8.25 |
| Train on internal images<br>Test on internal images | 62.77 | 6.78 | 21.93 | 9.14 | 3.67 | 9.50 | 0.81 | 22.34 |
| Train on internal/external images<br>Test on internal images | 59.79 | 6.46 | 20.18 | 8.41 | 2.84 | 7.56 | 0.96 | 17.46 |

As can be seen in the table, out-of-the-box, the Faster R-CNN models trained on our original rice images were not very accurate on the external images. In fact, the original models could not even detect the root in 4 out of 15 images, and could not detect the stele in 7 out of 15 images, due to the differences between the external images and our internal images used for training (if an object was not detected, a 0 diameter was assigned to it). However, the fine-tuned models significantly improved the results of the original models, with rRMSE dropping from 57.14% to 2.95% for RD, from 100.28% to 4.55% for SD, from 91.04% to 7.79% for LMXD, and from 53.96% to 8.25% for LMXN. We emphasize that the high errors of the original models are generally due to the models not being able to detect some objects at all (although the error for the objects detected was relatively small). These results show that the Faster R-CNN models fine-tuned with a small number of images (specifically, 42) can learn to predict the new types of images accurately.

To ensure that the performance of the fine-tuned models was not worse than the performance of the original models on our internal images, we also tested the fine-tuned models on the test fold corresponding to Split 1 (which was used for training). We recorded both the results of the original models and the results of the fine-tuned models in Table 4 (the last two rows, respectively). As can be seen, the results on our internal images improved slightly when using the fine-tuned models, as those models are more robust to variations. Specifically, rRMSE dropped from 6.78% to 6.46% for RD, from 9.14% to 8.41% for SD, from 9.50% to 7.56% for LMXD, and from 22.34% to 17.46% for LMXN. It is also interesting to note that the results of the models on the external images are better than the overall results on the internal images. One possible reason for this may be that the images found online are generally clearer images, used to illustrate root anatomy, despite the fact that they are different from our internal images.

### 3.4. Advantages and Disadvantages of the Faster R-CNN Approach

While a direct comparison between the Faster R-CNN model (trained on rice root cross-section images) and existing approaches (e.g., RootScan and RootAnalyzer) is not possible, given that each approach is trained on different species, in this section, we first outline several advantages of the Faster R-CNN model by comparison with existing models, and then emphasize several disadvantages.

Regarding the advantages, the following points can be made:

1. For an existing tool, it is hard to find parameters that are universally good for a set of images. For example, for a given set of parameters, the segmentation result from the RootAnalyzer in Figure 7 shows that the parameters are appropriate for the left rice image (a) where the LMX are reasonably well identified, but not appropriate for the right rice image (b) where no LMX is identified. As opposed to that, our experiments have shown that the performance of the Faster R-CNN model does not vary much with hyper-parameters. Once a model is properly trained, it performs accurately on a variety of images.
2. Plant samples used for imaging are grown in different conditions, for example in hydroponic (water based nutrient supply) or in soil, and root cross-section images are collected using different techniques (e.g., hand sectioning or sectioning using tools like vibratomes). Plant growing or image acquisition differences lead to differences in image’s color, contrast and brightness. As opposed to other tools, the Faster R-CNN model is not very sensitive to the light conditions or to the structure of the root cross-section images (including the epidermis thickness, epidermis transparency, and distorted cross-sections), assuming the models are trained with a variety of root cross-section images.
3. Each existing tool is designed with certain image characteristics in mind, and may not work on images that do not exhibit those characteristics. For example, RootAnalyzer assumes a clear cell boundary and does not work for images that contain a solid boundary where the cells are not clearly identifiable. The Faster R-CNN models simply reflect the broad characteristics of the images that they are trained on, instead of being built with some characteristics in mind. No specific background knowledge is provided, except for what is inferred automatically from training images.
4. Each tool is designed for a particular species, and incorporates back-ground knowledge for that particular species. As different species may have different root anatomy, a tool designed for a species may not work for other species. For example, RootAnalyzer is designed to automatically analyze maize and wheat root cross-section images, and “may work” for other species [48]. However, the Faster R-CNN model can be easily adapted to other species, assuming some annotated training images from those species are provided. No other background knowledge is required. Along the same lines, the Faster R-CNN model can be easily adapted to images with different resolutions, assuming those images include the features of interest.

**Figure 7:**
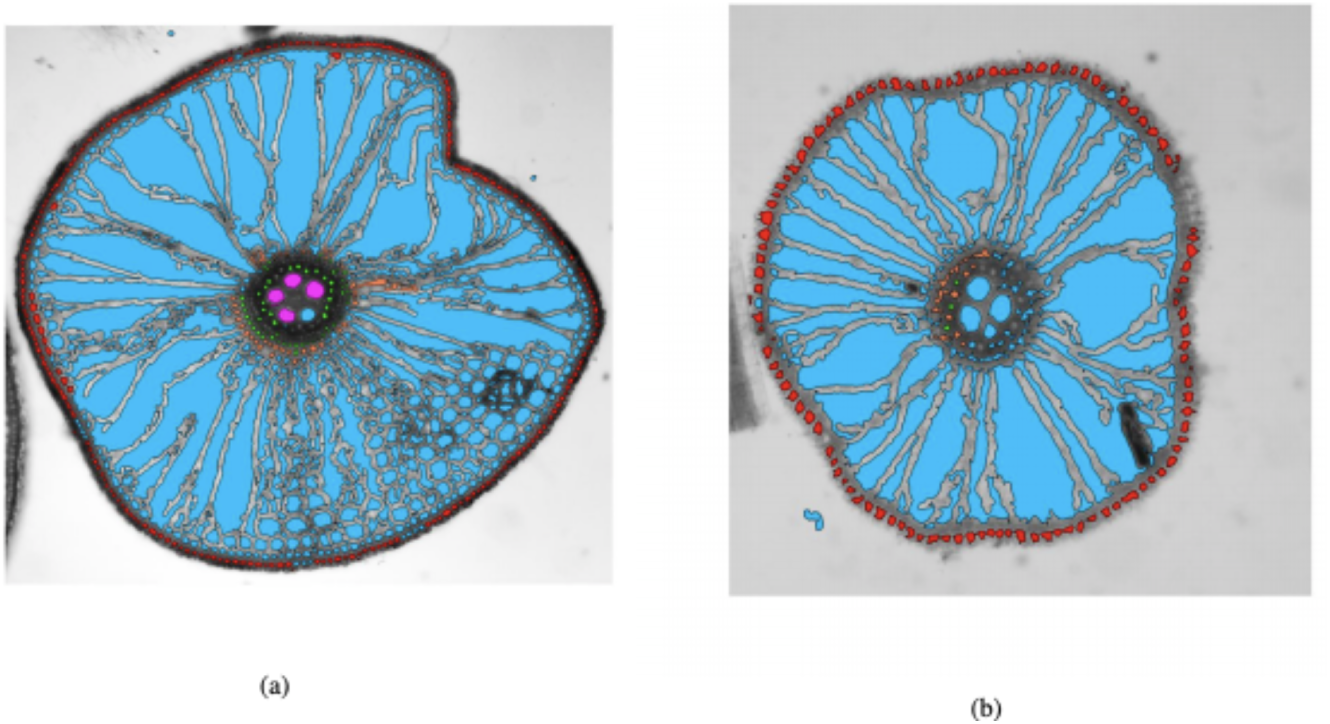
RootAnalyzer Annotations: With the same set of parameters, in the left image the root border (red), stele border (yellow), endodermis (green) and late-metaxylem (purple) are detected reasonably well, while in the right image, only half of the stele border is detected. Given that the tool fails to properly detect the stele border, it also fails to detect the late metaxylem.

While the Faster R-CNN model presents several advantages as compared to existing approaches that incorporate background knowledge, it also has several disadvantages, as outlined below:

- We found that smaller LMX objects are not detected by the Faster R-CNN models, most probably due to inconsistencies in the training data, as illustrated in Figure 6. To better handle noise and improve the performance, more training data might be needed. Alternatively, more consistent ground truth should be provided.
- While the bounding boxes which mark detected objects can produce accurate results, they are not always perfectly enclosing the detected object, as it can be seen in Figure 4. Thus, the diameter measurements can be sometimes slightly biased, and could potentially be improved.
- The Faster R-CNN model can accurately detect objects and identify traits such as diameter for the detected objects. The diameter can be subsequently used to derive other traits such as the object area. However, better area estimates could be potentially obtained with a Mask R-CNN model, which has the ability to detect object boundaries more precisely.
- The Faster R-CNN models can detect objects that can be marked with a bounding box. For other types of objects (e.g., aerenchym objects), Mask R-CNN models may be more appropriate.

### 3.5. Faster R-CNN Approach as a Tool for Root Anatomy

The Faster R-CNN model trained on our images can be used as a tool from a terminal or through a web-based application, which is also mobile friendly. The web-based application is available at https://rootanatomy.org. This site is linked to a GitHub repository that contains the source code, the pre-trained Faster R-CNN models and the ground truth data. The web-based application is user-friendly and does not require any programming skills. It can be run with one of our sample images displayed on the site, or with an image uploaded by the user.

### 3.6. Time Requirements

In terms of time/image requirements, our experiments have shown that accurate Faster R-CNN models can be trained from scratch with 150 to 250 images. The average time for labeling an image with LabelImg is approximately 2 minutes. The average time for training a model on an EC2 p2-xlarge instance available from Amazon Web Services (AWS) is approximately 10 hours, and does not require any human intervention during that time. Once the model is trained, the average time to annotate a new image is less than one second (using an EC2 p2-xlarge instance). If using our webserver (hosted on a local machine), the running time for annotating a new image is approximately 9 seconds, as this includes the time to setup the virtual environment, the time to retrieve the input image from the server, the time to perform the annotation, and the time to download the image to the user’s browser. Given these time requirements, assuming that a relatively large number of images need to be annotated for a biological study (on the order of thousands), the human time can be potentially reduced from days or weeks (the time would take to manually annotate all images) to hours (the time may take to manually label images for training) or minutes (the time for automatically annotating images with our tool).

Furthermore, the human time for labeling images for training could be dramatically reduced to less than an hour, if one is fine-tuning the Faster R-CNN model trained on our images as opposed to training a model from scratch.

## 4. Conclusions

In this paper, we trained Faster R-CNN models on rice root cross-section images and used the trained model to perform root anatomy. The Faster R-CNN approach to root anatomy is fully automated and does not need any background knowledge, except for the implicit knowledge in images that the model is trained on. More specifically, we trained Faster R-CNN models to detect root, stele and LMX objects, and to predict bounding boxes for each detected object. Subsequently, the bounding boxes were used to obtain anatomical properties, specifically, root diameter, stele diameter, LMX diameter and LMX number. The Faster R-CNN models used had VGG-16 as a backbone, to take advantage of the extensive training of the VGG-16 network, and were fine-tuned on root cross-section images.

We evaluated the Faster R-CNN models in terms of their ability to detect the objects of interest, and also in terms of their ability to lead to accurate measurements for RD, SD, LMXD and LMXN. The results of the evaluation showed that the models produced accurate and consistent annotations, when trained on a relatively small number of training images, specifically close to 300 images. For LMXD and LMXN, we trained Faster R-CNN models from both 50*×* magnification images and 100*×* magnification images. Our results showed that the performance is slightly better for the 100*×* magnification images, although this magnification is not a requirement for good performance. Furthermore, our results suggest that the Faster R-CNN models can potentially be used in practice to accelerate the speed at which root cross-section images are analyzed, and save significant human efforts and costs.

The evaluation in this paper was done on rice images. However, an important observation was that the models can be easily adapted to other types of root cross-section images and also to other species, by fine-tuning the existing Faster R-CNN models with a small number of labeled images from the species of interest. Similarly, additional anatomical features can be extracted by fine-tuning the existing models with images labeled according to other traits that are targeted (assuming the traits can be marked using bounding boxes).

While a direct comparison with existing tools for analyzing root cross-section images was not possible, we identified several advantages of the automated Faster R-CNN approach as compared to existing approaches that explicitly use background knowledge. We also identified several limitations of the Faster R-CNN model, including the fact that they can only be used for objects that can be represented using bounding boxes. As opposed to the Faster R-CNN model, existing approaches can identify a bigger variety of anatomical features. Thus, we can conclude that the Faster R-CNN approach and the existing tools have complementary strengths, and one cannot fully replace another.

As part of future work, we plan to thoroughly study domain adaptation approaches that allow the transfer of knowledge from the trained rice Faster R-CNN models to models for other plant species (or for other traits), without labeling a large number of images from the other species of interest.

## Author Contributions

XL carried out the original model adaptation and training, with input from DC. CW, XL and DC carried out the computational experiment design, with input from SVKJ. RB and SVKJ performed the biological experiment design and collection of the data. CW and XL carried out the computational experiments. RB performed the labeling of the data according to RD, SD, LMXD and LMXN measurements. CW and XL performed the bounding box labeling. SVKJ is the agronomy project leader with technical background in root phenotyping. DC is the computational project leader, with background in machine learning and deep learning. CW and XL drafted the first version of the manuscript, and DC and SVKJ contributed to the preparation of the final version of the manuscript. RJ contributed biological knowledge to the manuscript and provided feedback on the preliminary version. CW designed and developed the webserver. All authors read and approved the final manuscript.

## Data Availability Statement

The image datasets used in this study can be found in a GitHub repository at https://github.com/cwang16/Root-Anatomy-Using-Faster-RCNN.

## Acknowledgments

An earlier version of this manuscript has been released as a Pre-Print at https://www.biorxiv.org/content/10.1101/442244v2.article-info [65].

